# Inhibition of Early Response Genes Prevents Changes in Global Joint Metabolomic Profiles in Mouse Post-Traumatic Osteoarthritis

**DOI:** 10.1101/379370

**Authors:** Dominik R. Haudenschild, Alyssa K. Carlson, Donald L. Zignego, Jasper H.N. Yik, Jonathan K. Hilmer, Ronald K. June

## Abstract

Osteoarthritis (OA) is the most common degenerative joint disease, and joint injury increases the risk of OA by 10-fold. Although the injury event itself damages joint tissues, a substantial amount of secondary damage is mediated by the cellular responses to the injury. Cellular responses include the production and activation of proteases (MMPs, ADAMTSs, Cathepsins), the production of inflammatory cytokines, and we hypothesize, changes to the joint metabolome. The trajectory of cellular responses is driven by the transcriptional activation of early response genes, which requires Cdk9-dependent RNA Polymerase II phosphorylation. Flavopiridol is a potent and selective inhibitor of Cdk9 kinase activity, which prevents the transcriptional activation of early response genes. To model post-traumatic osteoarthritis, we subjected mice to non-invasive ACL-rupture joint injury. Following injury, mice were treated with flavopiridol to inhibit Cdk9-dependent transcriptional activation, or vehicle control. Global joint metabolomics were analyzed 1 hour after injury. We found that injury induced metabolomic changes, including increases in Vitamin D3 metabolism and others. Importantly, we found that inhibition of primary response gene activation at the time of injury largely prevented the global changes in the metabolomics profiles. Cluster analysis of joint metabolomes identified groups of injury-induced and drug-responsive metabolites, which may offer novel targets for cell-mediated secondary joint damage. Metabolomic profiling provides an instantaneous snapshot of biochemical activity representing cellular responses, and these data demonstrate the potential for inhibition of early response genes to alter the trajectory of cell-mediated degenerative changes following joint injury.

**Significance Statement:** Joint injury is an excellent predictor of future osteoarthritis. It is increasingly apparent that the acute cellular responses to injury contribute to the initiation and pathogenesis of OA. Although changes to the joint transcriptome have been extensively studied in the context of joint injury, little is known about changes to small-molecule metabolites. Here we use a non-invasive ACL rupture model of joint injury in mice to identify injury-induced changes to the global metabolomic profiles. In one experimental group we prevented the activation of primary response gene transcription using the Cdk9 inhibitor flavopiridol. Through this comparison, we identified two sets of metabolites that change acutely after joint injury: those that require transcription of primary response genes, and those that do not.

---

Osteoarthritis (OA) is the most common degenerative joint disease, occurring in >50% of the population older than age 65 (1). Changes in osteoarthritic joints include cartilage deterioration and degradation (2), low grade inflammation (3), fibrosis (4), subchondral bone remodeling (5), and other phenomena which comprise a complex etiology. At least 12% of all OA is directly caused by joint injury, and tears of the Anterior Cruciate Ligament (ACL) are the second most common knee injury (6–8). Within 10-20 years, up to 50% of patients with ACL tears develop radiographic evidence of OA (9, 10), representing a 10-fold increase in disease risk.

The increased risk for developing OA after injury results through multiple mechanisms. First, the mechanical damage that occurs during a joint injury has an immediate effect on the joint tissues: cell death and physical damage to joint tissues occurs within milliseconds of impact. Second, the immediate mechanical damage triggers an acute cellular response within minutes to hours(11). It is increasingly evident that the acute cellular response phase contributes substantially to the risk of developing arthritis after injury. The acute response phase is characterized by the release of inflammatory mediators such as IL1, IL6, iNOS, and TNF from the joint tissues(12). This, in turn, causes the transactivation of primary response genes and leads to the production of matrix-degrading enzymes such as MMPS, Cathepsins, Collagenases, Aggrecanases, and the production of inflammatory cytokines. The transactivation of the primary response genes thus constitutes a secondary wave of cell-mediated damage, which contributes substantially to the elevated risk for OA by causing irreversible damage at the molecular level(13).

The transcriptional activation of primary response genes is regulated by the kinase activity of cyclin-dependent kinase 9 (Cdk9). The majority of primary response genes are ‘primed’ for rapid transcriptional activation, with the transcription complex already assembled on the DNA promoters and the RNA polymerase complex stalled just before entering the transcriptional elongation stage. The rate-limiting step for the transcription of primary response genes is the recruitment of Cdk9, which then phosphorylates RNA Polymerase II to enable transcription to proceed. This central mechanism of regulation is highly conserved amongst primary response genes(14–18). Importantly, because RNA Polymerase II itself is the target for Cdk9 kinase, this regulatory mechanism is largely independent of the actual primary response gene being transcribed(18–20). Furthermore, multiple signaling pathways converge on Cdk9 kinase activity to initiate primary response gene transactivation. Flavopiridol, which is in phase II clinical trials as an anti-leukemia treatment (21), is one of several small-molecule inhibitors of Cdk9 with well-established pharmacokinetics. Flavopiridol inhibits Cdk9 by competing with adenosine triphosphate (ATP) for binding to the kinase active site (22). Previous research demonstrates that in vitro inhibition of Cdk9 using flavopiridol attenuates the transactivation of primary response gene transcription and protects chondrocytes and cartilage explants from apoptosis and cytokine-induced degradation (17).

One possible stimulus for the transactivation of primary response genes are the injury-induced changes in small molecule metabolites. Conversely, changes in the profile of transcribed genes can alter metabolism and thereby affect the relative abundance of small molecule metabolites. The abundance of small-molecule metabolites can be assessed quantitatively through metabolomic profiling. Metabolomic profiling describes the instantaneous cellular response through quantitative measurements of biochemical mediators (23). We hypothesized that the joint metabolome would exhibit global alterations during the acute post-injury time-frame. We further hypothesized that treatment with the Cdk9 inhibitor flavopiridol would partially block injury-induced changes in the joint metabolome by preventing activation of primary response genes.

To address our hypotheses, we profiled large-scale molecular changes in joint biology one hour following injury, in mice treated with the Cdk9 inhibitor flavopiridol or vehicle controls. We used two distinct metabolomics approaches: First, untargeted metabolomic profiling provides the opportunity for discovery of both novel and previously-defined biochemical mediators of the cellular response. Second, targeted metabolomic analysis allows for the sensitive quantification of small molecule expression levels (*e.g*. co-factors, hormones, vitamins, etc.), which describe known biological networks such as central energy metabolism (23). We observed that joint injury altered the joint metabolome at the one hour time-point. Many of the injury-induced metabolomic changes were attenuated in mice treated with Cdk9 inhibitor, suggesting that transcription of primary response genes is required for the changes to these metabolites.

## Results

### Joint Injury Altered Global Metabolomic Profiles

The right knee (stifle) joints of mice were injured by application of a mechanical load to the ankle, causing anterior translation of the tibia and stretching the anterior cruciate ligament (ACL) beyond the point of failure (Figure 4). Left knees were not injured and served as contralateral uninjured control joints. The entire injury takes under 2 seconds, and since this injury is a non-invasive method, it enables the study of the natural progression of the acute injury responses, including changes in gene expression and metabolite concentrations.

To assess functional changes in molecular biology related to joint injury, metabolites were extracted from micro-dissected joint tissue following injury. For injured mice, we found distinct metabolomic profiles between the injured right joints and contralateral uninjured left joints (Figure 1A). There were 66 metabolites detected in the injured knees that were not detected in the contralateral uninjured joints, and 81 metabolites detected in the contralateral uninjured joints that were undetected in the injured joints (Figure 1B). Overall, there were 132 metabolites upregulated and 131 metabolites down-regulated in injured joints compared with the contralateral uninjured joints of the same mice. Two clusters of interest were identified in hierarchical cluster analysis (HCA), with cluster 1 containing 35 metabolites downregulated with injury and cluster 2 with 47 metabolites upregulated with injury (Figure 1A). Within these clusters, we identified several metabolites differentially regulated by injury, including upregulation of anandamide and downregulation of glutamine (Figure 1C-D). We found substantial upregulation of metabolites related to vitamin D3 signaling in injured joints (Figure 1D). We observed downregulation of deoxycytidine triphosphate consistent with injury-induced upregulation of primary response genes (Figure 1D). Enrichment analysis revealed that glutamine and glutamate, arginine and proline, and pyrimidine metabolism were downregulated after injury. Pathways upregulated after injury included retinoid metabolism, phospholipid biosynthesis, hydroxyproline degradation, and anandamide metabolism.

**Fig. 1.**
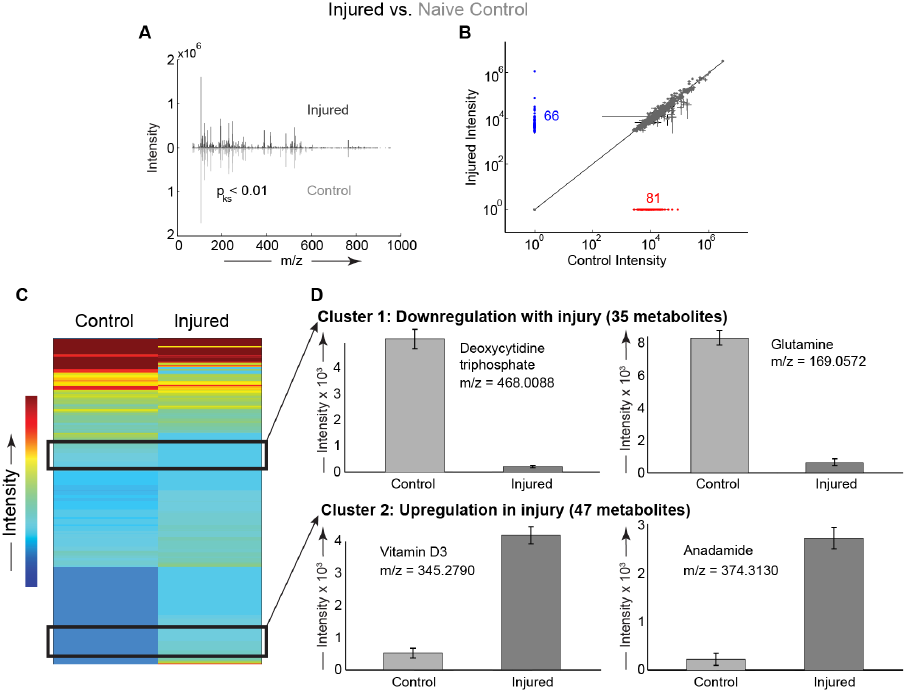
Joint injury induced global changes in the metabolomic profiles compared to contralateral uninjured joints of the same mice. (A) Distributions of metabolites differ between the control and injured knees (statistical comparison #2, see methods). Mirrored distributions plotted as median mass spectra from n = 6 mice. Injured knee plotted on top in black and uninjured knee plotted on bottom in gray. (B) Scatter plot comparing metabolite expression levels. There were 81 metabolites exclusively detected in uninjured control knees (red) and 66 metabolites exclusively detected in injured knees (blue). (C) Clustered heatmap of median metabolites between control and injured mouse knees. There were 822 metabolites detected in both control and injured joints. We identified two clusters of interest. (D) Cluster 1 includes 35 metabolites with decreased expression in injury, including glutamine. Cluster 2 contains 47 metabolites with increased expression in injured knees compared with uninjured knees. Cluster 2 includes a vitamin D3 derivative and anandamide. For panel D, all metabolites significantly different with all p < 0.01 using statistical comparison #2.

### Cdk9 Inhibition Prevents Injury-induced Metabolic Changes

Following injury, a group of mice was treated with the Cdk9 inhibitor flavopiridol to downregulate transcription of early response genes. Early response genes are associated with inflammation and degradative enzymes, which can lead to long-term joint damage. We observed distinct metabolomic profiles between joints of injured mice and joints of injured mice treated with flavopiridol (FP, Figure 2). The metabolomic profiles of injured joints treated with FP were more similar to control joints than to injured joints without treatment. In injured joints, there were 56 metabolites detected after FP-treatment that were not detected in untreated joints. In contrast, in injured joints there were 98 metabolites detected in untreated joints that were not present after FP-treatment. Treatment with FP abrogated the injury-induced upregulation of vitamin D3, phylloquinone, and acetylcarnitine. Enrichment analysis of the metabolite cluster upregulated after injury identified ubiquinone biosynthesis, vitamin digestion and absorption, transport of fatty acids, and various lipid metabolism pathways. Flavopiridol prevented injury-induced decreases in the expression of several metabolites (Figure 2B).

**Fig. 2.**
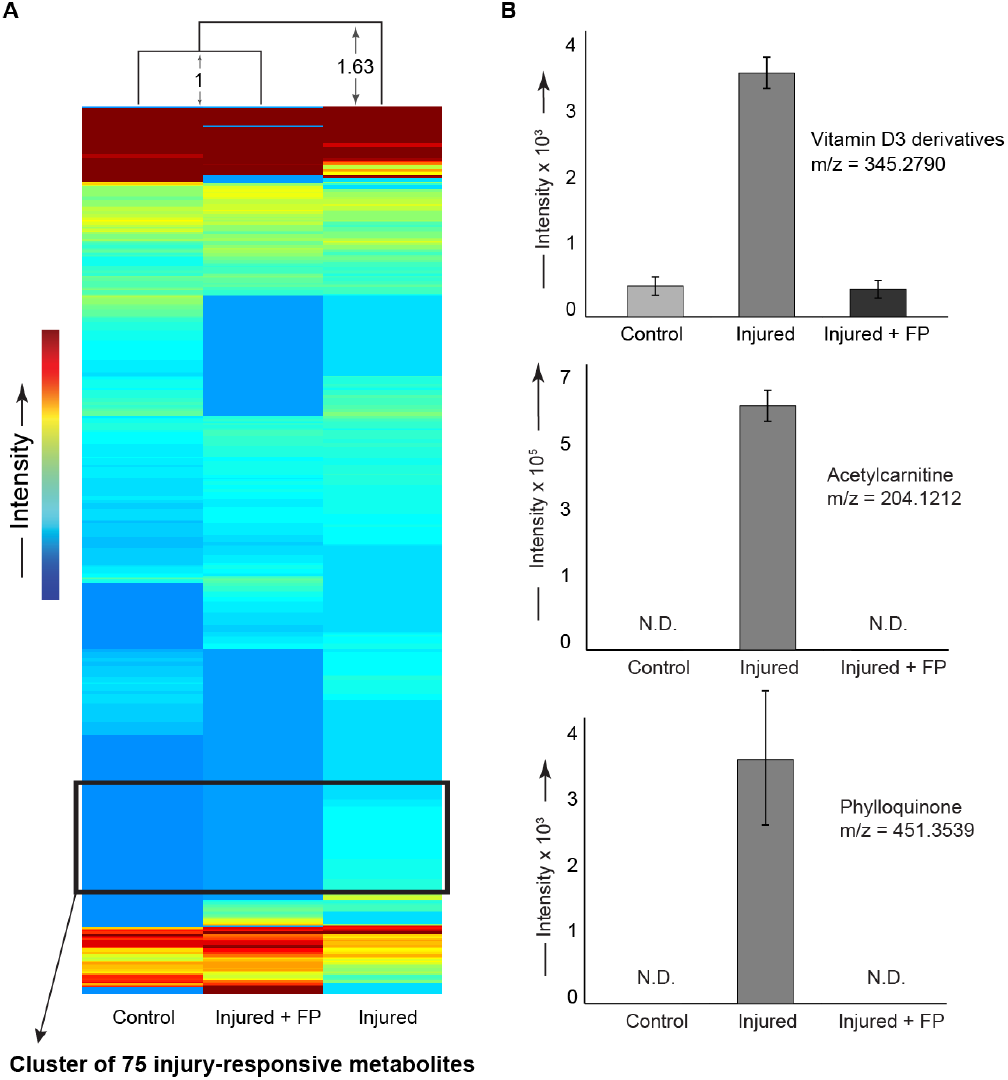
Intraperitoneal flavopiridol treatment after injury prevented global metabolomic changes induced by joint injury. All panels show right legs of mice subjected to either injury, injury and flavopiridol (FP), or uninjured controls from naïve mice that received neither injury nor drug, (statistical comparisons #4 and #6, see methods). (A) Unsupervised clustering performed on median values of 426 metabolites common to all experimental groups. The injured + flavopiridol group clustered most closely to the naïve uninjured control group. The distribution of injured mice was ~63% further than the distance between the control and flavopiridol-treated mice. We identified a cluster of 75 injury-responsive metabolites. (B) Selected metabolites upregulated by injury and rescued by flavopiridol treatment. Flavopiridol prevented the injury-induced increases in vitamin D3 and derivatives, acetylcarnitine, and phylloquinone. All p < 0.01 when comparing injured to either controls or injured + flavopiridol group using FDR-corrected t-tests for comparisons #4 and #6.

### Cdk9 Inhibition Represses Injury-Induced Changes in Joint Energy Metabolism

Because both joint degradation and repair processes require substantial energy via ATP hydrolysis (24), we quantified metabolites targeted to central energy metabolism (*i.e*. glycolysis, the pentose phosphate pathway, and the TCA cycle). By examining the ratios of downstream to upstream metabolites, we inferred changes in energy utilization.

Within the pentose phosphate pathway, there are two reactions that convert NADP^+^ to NADPH. Injury upregulated the ratio of NADPH to NADP^+^, consistent with the transcription of early response genes (Figure 3B). Treatment with flavopiridol was able to reduce this injury-induced upregulation. Injury also upregulated the ratio of ATP to ADP indicating increased glycolysis (Figure 3C). Flavopiridol-treated mice had a higher ratio of ATP to ADP suggesting increased glycolytic metabolism upon inhibition of early response genes.

**Fig. 3.**
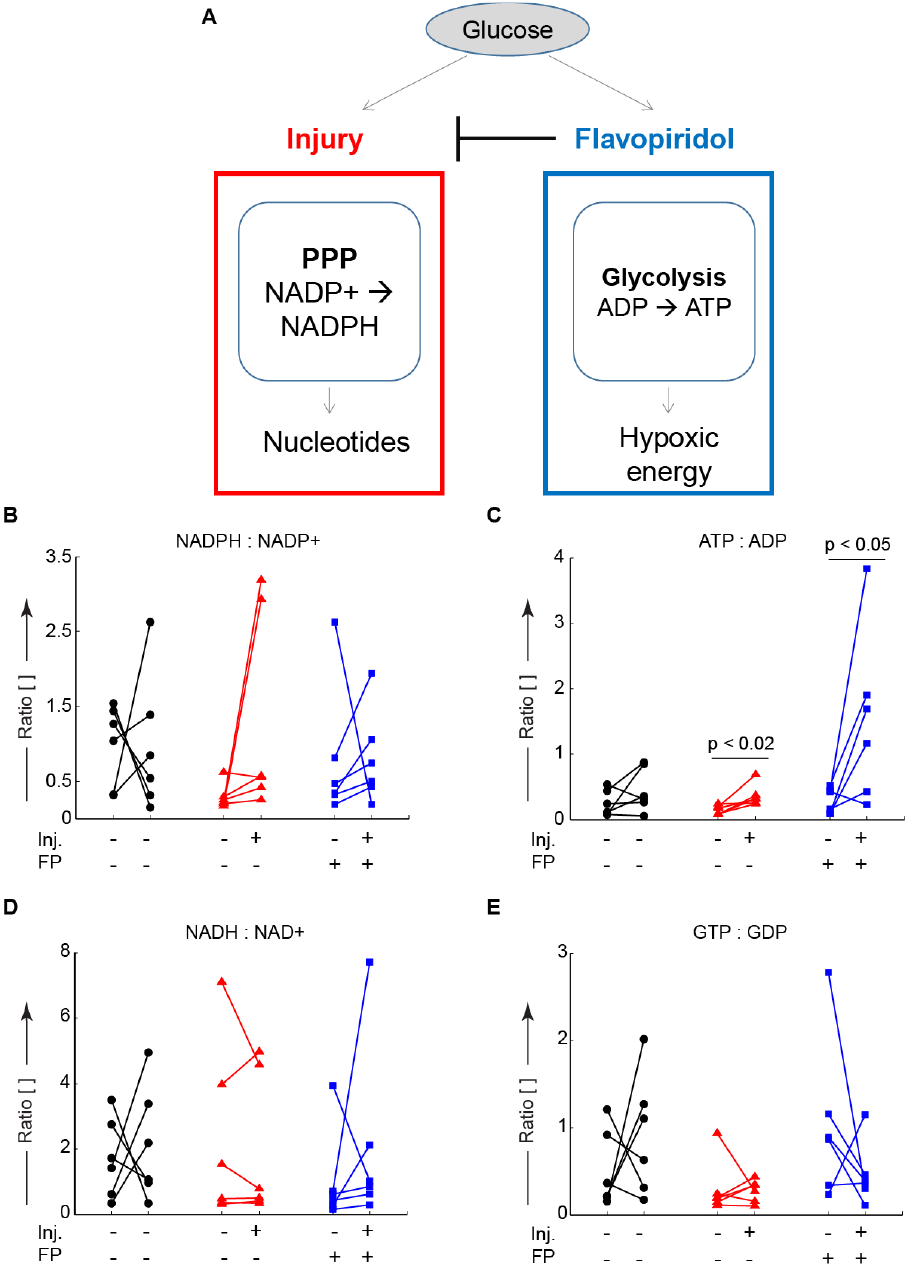
Glucose metabolism is altered by injury and partially rescued by Cdk9 inhibition. After joint injury and treatment with the Cdk9 inhibitor flavopiridol, the upstream to downstream ratios of central-energy-related metabolites were examined via LC-MS to understand energy utilization. (A) Conceptual model of glucose utilization injury upregulated pentose phosphate activity (PPP) to produce nucleotides. Flavopiridol blocks this upregulation and increases glycolytic metabolism. (A) There was a single outlier (plotted) in the NADPH:NADP^+^ ratio data. Five out of six mice demonstrated increases in the NADPH:NADP^+^ ratio upon indicating increased flux through the pentose phosphate pathway, potentially to produce nucleotides for upregulated gene expression. (B) The ATP:ADP ratio was increased in the injured knees for both control and treated groups indicating increased glycolytic flux, consistent with hypoxic energy utilization within the joint. (C) There were no significant changes in the ratio of NAD_+_:NADH. (D) The ratio of GDP:GTP was unchanged indicating similar TCA-cycle energy utilization. Data from n = 6 mice. In panels B-E, lines connect data from the injured and contralateral knees of the same mouse for paired analyses.

The ratio of NADP^+^ to NADPH was larger in the contralateral control joints than injured joints (Figure 3D), which was driven by increases in NADPH expression in the injured joint (Supplemental Figure S1). The ratio of ADP to ATP was increased in injured knees compared with contralateral controls, and this increase was accentuated via treatment with flavopiridol. The ratios of NAD+ to NADH and GDP to GTP were largely unchanged by injury. Taken together, these results indicate that injury decreases glycolytic metabolism with a concomitant increase in carbon metabolism via the pentose phosphate pathway, which may result in transcription of early response genes.

## Discussion and Conclusion

Flavopiridol treatment of mice after injury restored global metabolomic profiles to those of uninjured controls. Flavopiridol suppresses Cdk9-dependent phosphorylation of RNA Polymerase II, which is the rate-limiting step for the transcriptional elongation of primary response genes, and therefore effectively suppresses early response genes (17). In this study, we examined whether or not flavopiridol treatment would alter joint metabolomics following injury. Injured mice demonstrated global metabolomic changes, and therapeutic intervention with flavopiridol abrogated those changes demonstrating its potential for treating joint injuries in human patients.

Unsupervised clustering of 426 metabolites found the greatest similarity between the metabolomes of uninjured control joints from naïve mice and injured-FP-treated joints (Figure 2A). This suggests that activation of primary response genes within the first hour strongly contributes to the metabolomic response to injury.

Using high-dimensional metabolomics, we identified several novel pathways affected by joint injury. Previous studies have utilized metabolomics to assess chondrocyte mechanotrans-duction (25) and to compare normal and OA synovium (26). Here we apply metabolomic profiling to study acute changes upon joint injury in mouse knees. We find that joint injury alters both global and targeted metabolomic profiles.

In this model, vitamin D3 and derivatives and vitamin digestion and absorption pathways were upregulated after joint injury. This injury-induced upregulation was reversed by treatment with flavopiridol after injury, suggesting that the transcription of primary response genes is involved in mediating these changes. These observations are likely related to the changes in subchondral bone seen in both mouse and human OA(27). Increased vitamin D signaling has previously been reported in arthritic joints in response to joint damage to induce new bone formation, with this increased bone remodeling associated with joint pain(28, 29). In this noninvasive injury model, we observe dramatic bone remodeling as early as 3 days postinjury, resulting in a 25-35% loss of subchondral bone volume peaking at 7-10 days post-injury(30). Since bone is innervated, modulation of vitamin D3 signaling in OA may provide novel therapeutic strategies for improving osteoarthritis pain, which is the major cause of debilitation(31). While previous clinical trials have administered vitamin D3 to OA patients(32), these results suggest that inhibition of vitamin D3 signaling, as achieved with Cdk9 inhibition here, may be beneficial for OA patients. The data potentially show that injury induces vitamin D3 metabolism, which may be partially blocked by Cdk9 inhibition, indicating that blocking primary response gene transcription may minimize injury-induced changes in subchondral bone.

Injury increased levels of phylloquinone (vitamin K) in the joint, with this increase abrogated by flavopiridol treatment. Phylloquinone is the primary form of vitamin K, and sufficient vitamin K levels are imperative for normal cartilage and bone mineralization(33). Previous research found primary vitamin K levels are lower in chronic hand and knee OA(34). Interestingly, we found increased levels of phylloquinone (vitamin K) and vitamin digestion and absorption pathways upregulated after injury. These injury-induced increases are likely due to the bone remodeling caused by the mechanical overload. Furthermore, the flavopiridol-induced reduction in phylloquinone levels may further suggest that transcription of early response genes mediates bone remodeling after injury.

Acetylcarnitine was upregulated in the joint after injury. Acetylcarnitine is metabolized to carnitine, which transports fatty acids to the mitochondria for breakdown. We also identified fatty acid transport and various lipid metabolism pathways upregulated after injury. Lipids and fatty acids have previously been implicated in chondrocyte mechanotransduction, although their importance in OA pathogenesis remains unknown(35). Levels of acetylcarnitine were abrogated with flavopiridol treatment, which may suggest that transcription of early response genes may be mediating injury-induced perturbations in fatty acid metabolism. Previous studies have also investigated the prophylactic role of acetylcarnitine in a monosodium iodoacetate rat model of OA, demonstrating that acetylcarnitine supplementation was able to reduce overall OA damage in the joint and attenuate OA pain(36). Flavopiridol treatment, however, did abrogate the increase in acetylcarnitine levels in this study, which could suggest a potentially negative side effect of inhibiting the transcription of early response genes after injury. Further studies are needed to elucidate the role of fatty acid metabolism and acetylcarnitine levels after injury and its implications for the development of PTOA.

Anandamide levels and anandamide metabolism were up-regulated after joint injury. A likely source of anandamide in joints is synovial fibroblasts because these cells contain the required enzymes for its synthesis(37). Elevated levels of anandamide have previously been found in synovium and synovial fluid from patients with end-stage OA(38). Anandamide is an endocannabinoid implicated in inflammation, pain, nociceptive signaling, and analgesia via cannabinoid receptors. Importantly, elevated levels of anandamide after injury are thought to modulate the intensity of pain stimuli. Genes required for anandamide synthesis are induced by LPS within 90 minutes(39), but to our knowledge they have not been studied specifically as Cdk9-dependent primary response genes. Our results support increased anandamide levels after injury. However, additional studies are needed to elucidate anandamide’s role in injury-induced pain signaling. Anandamide also has known roles in osteoclast and osteoblast activation and proliferation(40, 41). Taken together with the subchondral bone remodeling after ACL rupture in this model, the increased levels of anandamide after injury may also suggest a role in injury-induced bone remodeling. Consistent with previous findings, these results suggest that elevated anan-damide levels are associated with injury and may be playing a role in injury-induced bone remodeling and/or pain signaling.

Targeted metabolomics found several changes in central energy metabolism following joint injury. The flux through the pentose phosphate pathway, as quantified by decreases in the ratio of NADP^+^ to NADPH increased after joint injury. This increase is likely due to the increased requirement for nucleobases to transcribe early response genes. As expected, this flux increase was abrogated following Cdk9 inhibition via flavopiridol treatment. Similarly, glycolytic flux, quantified by decreases in the ratio of ATP to ADP decreased following joint injury. We found no changes in the TCA cycle represented by the ratio of GTP to GDP. These two observations are consistent with the joint as a hypoxic environment where most energy utilization is via anaerobic glycolysis(42).

The ability of Cdk9 inhibitor flavopiridol to restore global changes in metabolomics caused by injury in mice is promising for improving treatment of injury-related osteoarthritis. Flavopiridol has been tested in human patients as a Cdk9 inhibitor for various cancers and is currently in phase II clinical trials as an anti-leukemia drug. Extension of this study will test the ability of flavopiridol to reduce cartilage deterioration and subchondral bone changes following mouse and human joint injury. In conclusion, this study demonstrated (1) the utility of high-dimensional metabolomic analysis to examine global changes in molecular biology in animal models and (2) the potential for Cdk9 inhibition as a strategy to improve joint health following injury.

## Materials and Methods

### Joint Injury Model and Experimental Design

Eighteen 12-week old C57BL/6 male mice were obtained from Jackson Labs, and randomly assigned into three experimental groups: 1) Uninjured, 2) Injured, and 3) Injured and Injected with Flavopiridol. Mice were anesthetized with isoflurane inhalation, and the right knees injured exactly as described (30). Briefly, the right ankle and knee joint was positioned in custom-machined platens in a mechanical testing setup (Figure 4). A single mechanical load was applied to the ankle (1mm/s to 12N) causing anterior translation of the tibia relative to the femur, stretching the ACL beyond the point of failure. The left knees served as uninjured contralateral controls. Immediately after injury, before waking from anesthesia, mice were given a systemic dose of 7.5mg/kg Flavopiridol by intra-peritoneal injection. Joints were harvested 1 hour after injury following sacrifice via CO2 inhalation. All superficial soft tissue (skin, muscle etc.) was removed. The joint was isolated via scalpel between the tibial and femoral growth plates, which included subchondral bone, synovium, articular cartilage, menisci, and ligaments. All mouse experiments were conducted in accordance with ethical standards and with approval from the institutional IACUC.

**Fig. 4.**
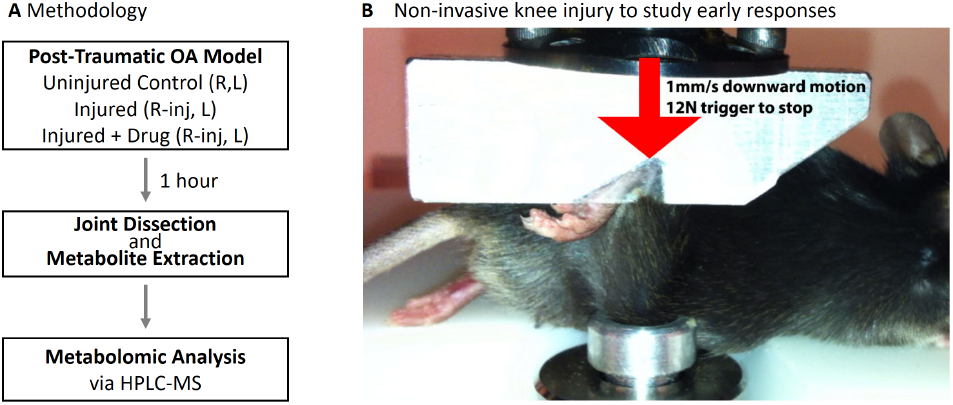
Non-invasive mouse ACL-rupture system to study acute injury responses in OA pathogenesis. (A) Experimental Design: One hour after injury, the individual metabolomes from joint tissues were characterized via LC-MS. (B) Mice were subjected to a single mechanical overload of 12 Newtons applied at 1 mm/s that results in ACL rupture to the right knee. This initiates a rapid induction of catabolic and inflammatory genes, and subsequent pathophysiological changes that mimic human osteoarthritis, including GAG loss and subchondral bone changes. The left knees serve as contralateral controls.

#### A. Metabolite Extraction

Metabolites were extracted using our previous methods (43), with modifications for joint tissue. Joint tissue was immediately frozen in liquid nitrogen and pulverized. Metabolites were extracted from pulverized joints in methanol:acetone (70:30) with repeated vortexing followed by extraction at −20°C overnight. Samples were pelleted at 4°C and 13,000 rpm for 10 minutes, the supernatant extracted, and the solvent removed via centrifugation under a vacuum for 8 hours at room temperature. Prior to running the dried samples on the mass spectrometer, samples were resuspended in 100 *μ*l of mass spectrometry grade water:acetonitrile (50:50 v/v).

#### B. Metabolomic Profiling

Metabolite detection was performed in the Montana State University Mass Spectrometry Core Facility via HPLC-MS using previously optimized protocols(25, 35). Chromatography was performed using an aqueous normal-phase, hydrophilic interaction chromatography (ANP/HILIC) HPLC column (Cogent Diamond Hydride Type-C) with a length of 150 mm, diameter of 2.1 mm, and particle size of 4 *μ*m in diameter. The column was coupled to an Agilent 1290 HPLC system, and chromatography used previously optimized methods (43). Following each run, blank solvent samples were run to ensure thorough washing of the column.

#### C. Data Analysis and Statistics

Data analysis involved both metabolites targeted to central energy metabolism and global untargeted metabolites. For the untargeted approach, raw HPLC-MS data were converted into .mzXML files using MS Convert (ProteoWizard, (44)), and processed in MZmine2.0 (45) prior to statistical analysis. In MZmine2.0 the datasets were filtered using established methods (43, 45) as follows: chromatograms were created using a minimum signal level of 1000 m/z, m/z tolerance of 15 ppm and a minimum time span of 0.1 minutes. Chromatograms were then normalized using a minimum standard intensity of 1000 m/z, at a 15 ppm tolerance, and a 0.25 min retention time tolerance. Chromatograms were then aligned using a 15 ppm tolerance for both retention time and mass. Following chromatogram alignment, lists were created and used for statistical analysis and metabolite identification.

For the targeted approach, ~50 metabolites known to be involved in central energy metabolism (46, 47) were analyzed. MassHunter’s Quantitative Analysis package (Agilent Technologies) was used to create a list of the calculated isotopic distributions (H^+^ and Na^+^ adducts) of the ~50 specific targeted masses (Isotope Distribution Calculator, Agilent Technologies). For putative metabolite identification, retention times were used with matched values to those from standard analytical samples determined and maintained by the MSU Mass Spectrometry Core. For each of the targeted metabolites, a 20 ppm window was set for each m/z value.

To assess the effects of Cdk9 inhibition on global joint biology following traumatic joint-injury, six separate sample groups were established, each with sample size n = 6 mice: Uninjured/control mice left leg (CL), uninjured/control mice right leg (CR), injured mice left leg (IL), injured mice right leg (IR), injured + flavopiridol mice left leg (FIL), and injured + flavopiridol mice right leg (FIR) (Table I). The left and right legs of the uninjured/control mice were not subjected to injury. For the injured mice, the right leg was subjected to joint injury, whereas the left leg served as a sham control, and was not subjected to injury. For the injured + flavopiridol mice, the right leg was subjected to injury and the left leg served as a sham control and was not subjected to injury. The flavopiridol was administered systemically by IP injection. For all statistical analysis, we defined detected masses as those present in the majority of samples.

**Table 1.** Designation of Experimental Groups

| Control Uninjured<br>(n=6) | Injured<br>(n=6) | Injured+Flavopiridol<br>(n=6) |
| --- | --- | --- |
| CR (Control Right) | IR (Injured Right) | FIR (FP-treated Injured Right) |
| CL (Control Left) | IL (Contralateral Left) | FIL (FP-treated Contralateral Left) |

Global metabolomic profiles were visualized by hierarchical agglomerative cluster analysis (clustergram in MATLAB) of the median profile of each group. Clusters of co-regulated metabolites were identified. Six statistical comparisons were made in analyzing the data: (1) CL vs. CR, (2) IL vs. IR, (3) FIL vs. FIR, (4), CR vs. IR, (5) CR vs. FIR, and (6) IR vs. FIR. To find statistically significant differences, t-tests were used (48) with significant differences identified by p-values less than 0.05. To minimize the risk of false positives associated with multiple comparisons testing, standard false discovery rate (FDR) calculations (48) were used with a q-value of 0.05.

To assess the differences in the metabolite intensity distributions (m/z spectra plots for the various sample groups), two-sample Kolmogorov-Smirnov tests were used, with statistically significant differences identified by p≤0.05. Kolmogorov-Smirnov tests are used as a non-parametric test for the equality of distributions between two samples. The same six comparisons were made: (1) CL vs. CR, (2) IL vs. IR, (3) FIL vs. FIR, (4), CR vs. IR, (5) CR vs. FIR, and (6) IR vs. FIR. Targeted metabolite profiles were analyzed by hierarchical agglomerative cluster analysis and the median ratios of NADP^+^:NADPH, NAD^+^:NADH, ATP:ADP, and GDP:GTP were calculated for each of the sample groups to assess relative changes in energy metabolism.

### Compound Identification and Pathway Enrichment

For putative metabolite identification in the untargeted metabolomic approach, a batch search of all of the metabolite mass to charge (m/z) values was performed in METLIN and HMDB whose databases contain over 80,000 identifiable metabolites (49, 50), including lipid identifications from LipidMAPS (51). Search parameters included using a mass tolerance of 20 ppm, and positively charged molecules with potential +1H^+^ or +1Na^+^ adducts. Putative metabolite identities within clusters of interest identified by hyerarchical cluster analysis were assessed for enriched pathways in IMPaLA(52).

## ACKNOWLEDGMENTS

We would like to acknowledge the Mass Spectrometry CORE at Montana State University for their technical assistance in mass spectrometry. This work was supported in part by CDMRP/PRMRP Grant PR110507 to DRH and by NSF CMMI 1554708 to RKJ. The funders had no role in study design, data collection and analysis, decision to publish, or preparation of the manuscript.

